## Supplemental Figures for "Prefoldin 5 is a microtubule-associated protein that suppresses Tau-aggregation and neurotoxicity"

**Supplementary Information**

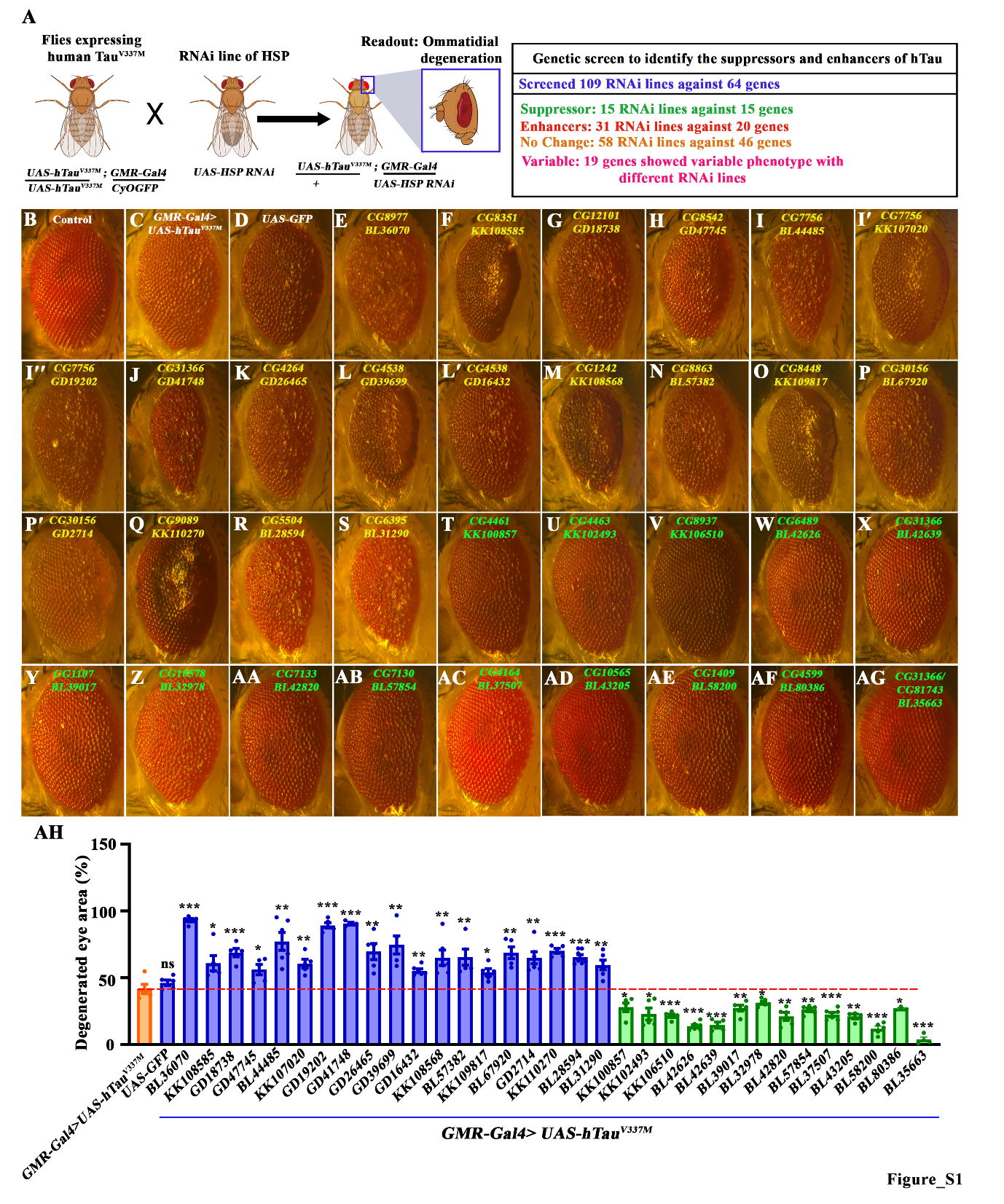

**Figure S1: Screening of *Drosophila* chaperones to identify modifiers of Tauopathy**

**(A)** Schematic representation of the genetic screen performed to identify the chaperones that function as modifiers of Tauopathy. In this screen, 109 RNAi lines corresponding to 64 chaperones were used, among which 15 RNAi lines against 15 genes were scored for their role as suppressors. While 20 genes (31 RNAi lines) enhanced the hTau phenotype, 46 genes (58 RNAi lines) did not alter the hTau phenotype in the eyes. Nineteen genes showed variable results with different RNAi lines, which we did not consider as either enhancers or suppressors.

**(B-AG)** Brightfield images of 7-day-old *Drosophila* eyes coexpressing *hTau^V337M^* and various RNAi lines against *Drosophila* chaperones.

**(AH)** Histogram showing the percentage of degenerated area in 7-day flies in coexpressing *hTau^V337M^* and various RNAi lines against chaperones. At least 3 brightfield eye images of each genotype were used for quantification. The details of the RNAi lines and statistical analysis are listed in Table 1.

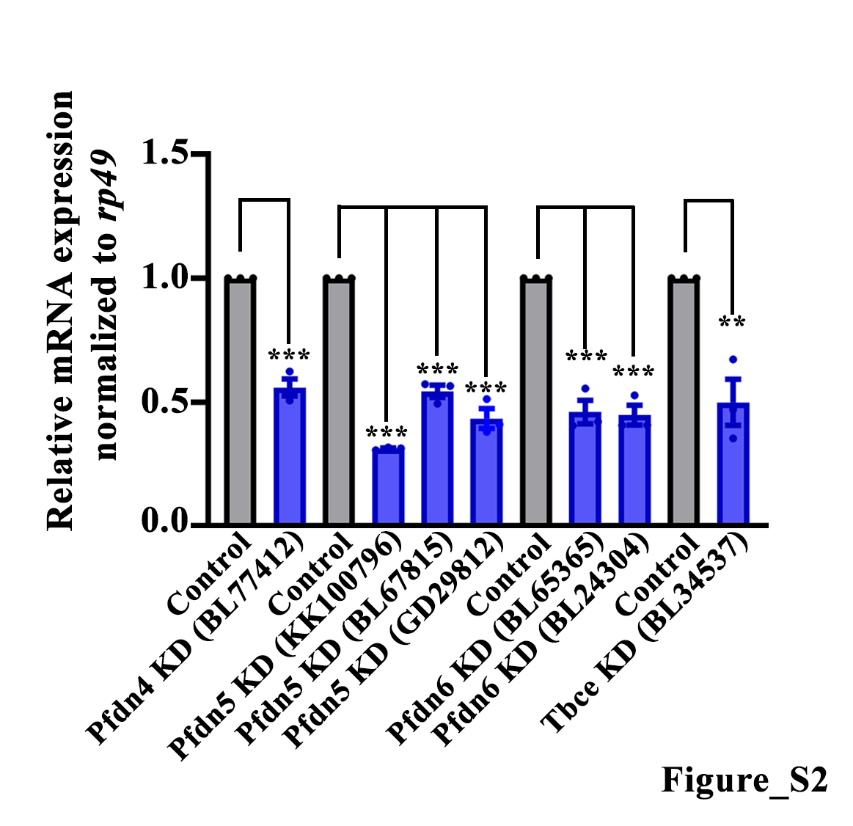

**Figure S2:** **qPCR analysis for knockdown efficiency of RNAi lines against cytoskeleton regulatory chaperones.**

Histogram showing the transcript levels of the mentioned genes. Total RNA was isolated from larval fillets of Actin-Gal4 driven RNAi lines. *rp49* was used as an internal control. The knockdown animals showed about 50-60% reduction in the transcript level. Three independent qRT-PCRs were performed for each genotype. Error bars represent SEM. ***p<0.001, **p<0.01.

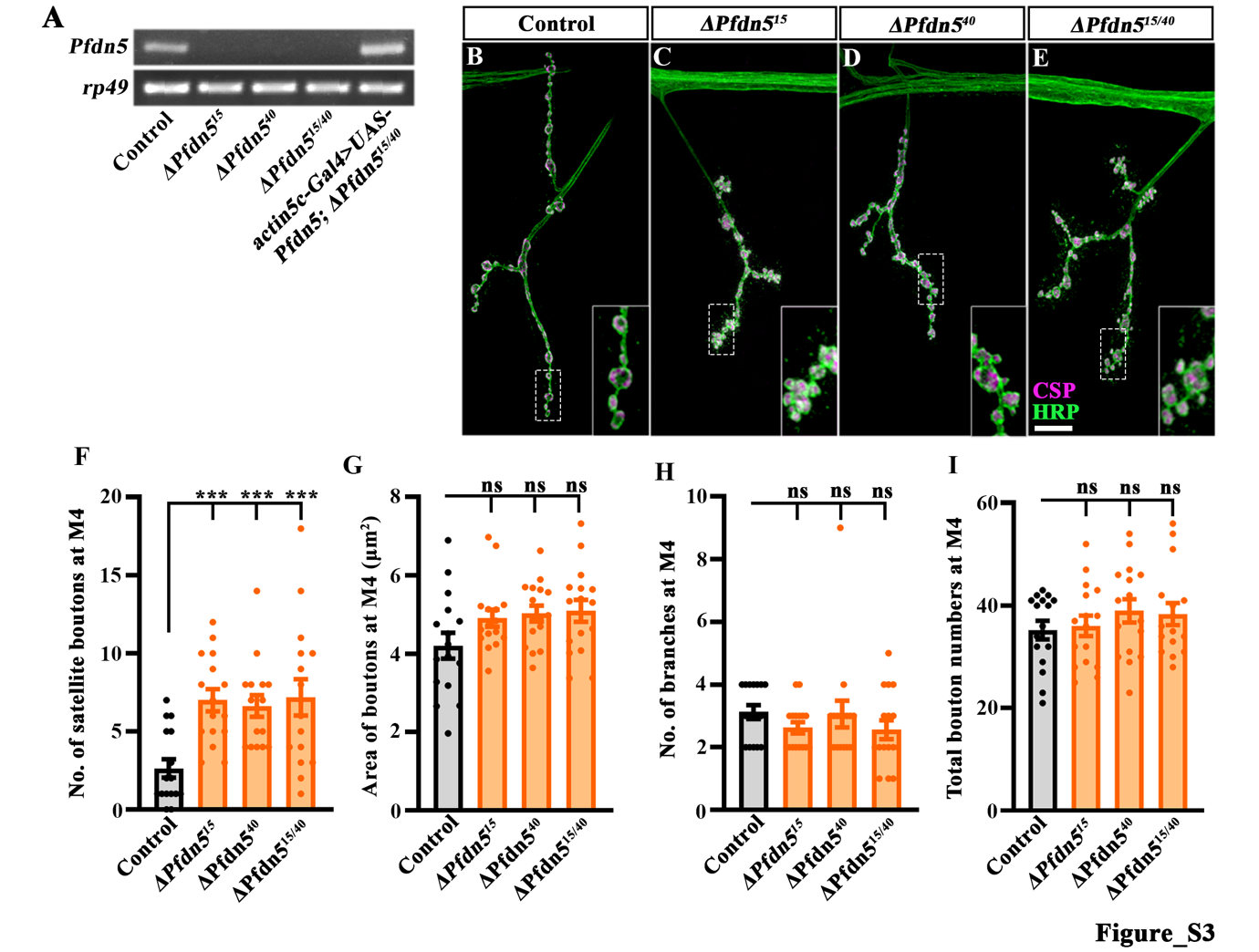

**Figure S3: *Pfdn5* mutants are null alleles with no detectable *Pfdn5* transcripts**

**(A)** Semi-quantitative RT-PCR showing transcript levels of *Pfdn5* in control, *ΔPfdn5^15/15^*, *ΔPfdn5^40/40^*, *ΔPfdn5^15/40^*, and *actin5C-Gal4>UAS-Pfdn5; ∆Pfdn5^15/40^*. *rp49* was used as an internal control for RT-PCR.

**(B-E)** Confocal images of NMJ synapses at muscle 4 of A2 hemisegment showing synaptic morphology in (B) control, (C) *ΔPfdn5^15/15^* (D) *ΔPfdn5^40/40^* (E) *ΔPfdn5^15/40^* double immunolabeled for HRP (green) and CSP (magenta). The insets represent the 3X magnified portion of the image shown in a white box. The scale bar in E for (B-E) represents 10 µm.

**(F)** Histogram showing the number of satellite boutons from muscle 4 at A2 hemisegment in control (2.63 ± 0.59), *ΔPfdn5^15/15^* (7.0 ± 0.7)*, ΔPfdn5^40/40^* (6.63 ± 0.7)*,* and *ΔPfdn5^15/40^* (7.2 ± 1.1). ***p<0.001. At least 16 NMJs of each genotype were used for quantification.

**(G)** Histogram showing bouton area (in µm^2^) from muscle 4 at A2 hemisegment in control (4.2 ± 0.3), *ΔPfdn5^15/15^* (4.9 ± 0.2)*, ΔPfdn5^40/40^* (5.0 ± 0.2)*,* and *ΔPfdn5^15/40^* (5.1 ± 0.3). ns, not significant. At least 16 NMJs of each genotype were used for quantification.

**(H)** Histogram showing the number of branches per NMJ from muscle 4 at A2 hemisegment in control (3.12 ± 0.2), *ΔPfdn5^15/15^* (2.6 ± 0.2)*, ΔPfdn5^40/40^* (3.10 ± 0.4)*,* and *ΔPfdn5^15/40^* (2.6 ± 0.3). ns, not significant. At least 16 NMJs of each genotype were used for quantification.

**(I)** Histogram showing total bouton number from muscle 4 at A2 hemi segment in control (35.25 ± 1.8), *ΔPfdn5^15/15^* (36.06 ± 2.0)*, ΔPfdn5^40/40^* (39.0 ± 2.3)*,* and *ΔPfdn5^15/40^* (38.4 ± 2.1). ns, not significant. At least 16 NMJs of each genotype were used for quantification.

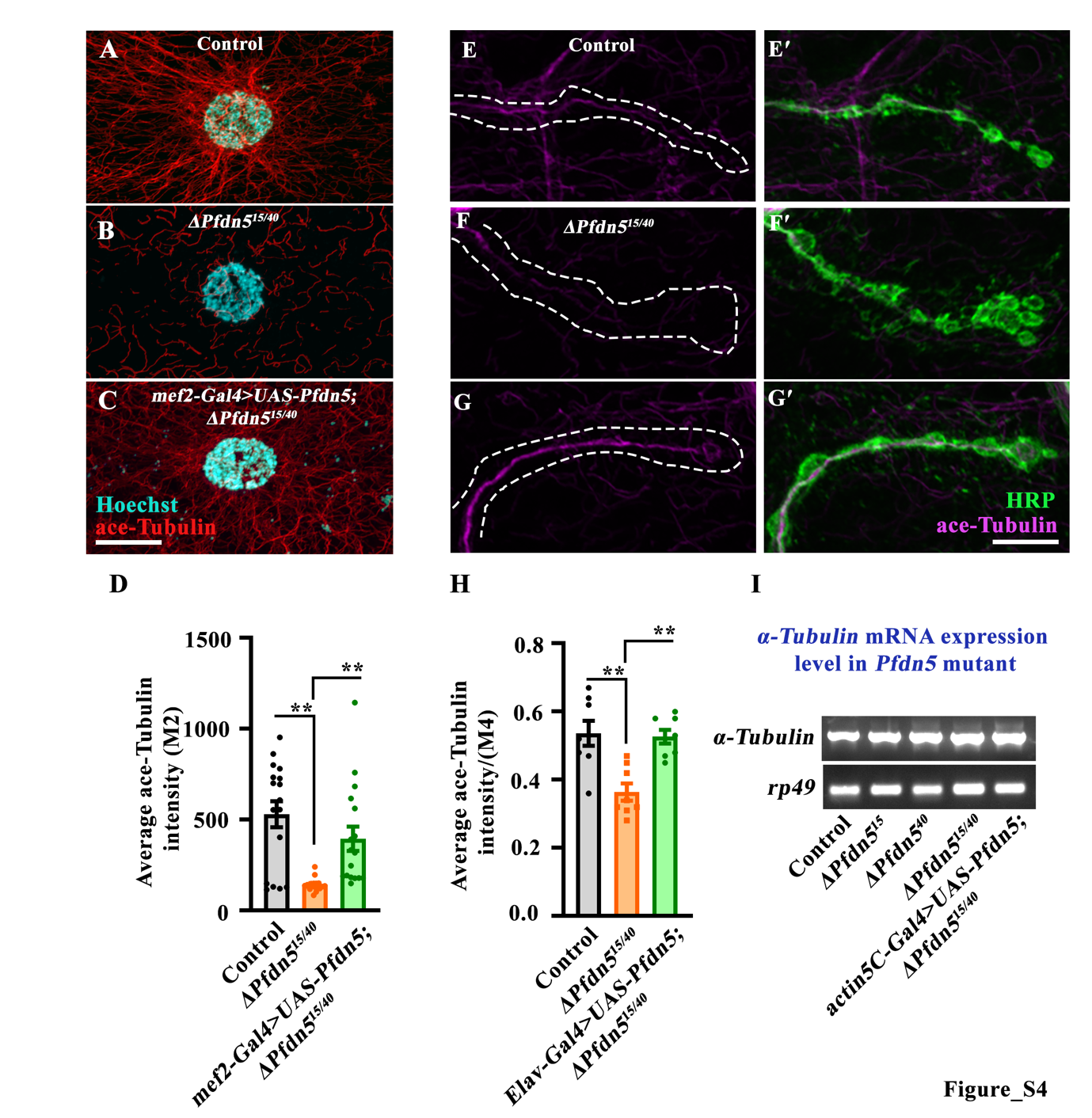

**Figure S4: Loss of Pfdn5 disrupts microtubule cytoskeleton**

**(A-C)** Confocal images showing the muscle 2 of A2 hemisegment in control, *ΔPfdn5^15/40^,* and *mef2-Gal4>UAS-Pfdn5; ΔPfdn5^15/40^* double immunolabelled with ace-Tubulin (red) and Hoechst (cyan). The inset shows the magnified version of the muscle nucleus and microtubule network. The scale bar in the C represents 20 µm for images (A-C).

**(D)** Histogram showing the average fluorescence intensity (au) of ace-tubulin at muscle 2 of A2 hemisegment in control (530.1 ± 71.6), *ΔPfdn5^15/40^* (141.9 ± 8.43)*,* and *mef2-Gal4>UAS-Pfdn5; ΔPfdn5^15/40^* (395.1 ± 66.78). **p<0.01. At least 17 NMJs of each genotype were used for quantification.

**(E-G')** Confocal images of NMJ synapses at muscle 4 of A2 hemisegment showing synaptic levels of ace-tubulin in control, *ΔPfdn5^15/40^,* and *Elav-Gal4>UAS-Pfdn5; ΔPfdn5^15/40^* double immunolabelled for ace-Tubulin (magenta) and HRP (green). The scale bar in the G' represents 20 µm for images (E-G').

**(H)** Histogram showing average fluorescence intensity of ace-tubulin at muscle 4 NMJ in control (0.54 ± 0.04), *ΔPfdn5^15/40^* (0.36 ± 0.03)*,* and *Elav-Gal4>UAS-Pfdn5; ΔPfdn5^15/40^* (0.53 ± 0.02). **p<0.01. At least 8 NMJs of each genotype were used for quantification.

**(I)** Semi-quantitative RT-PCR showing transcript levels of α-tubulin in control, *ΔPfdn5^15/15^, ΔPfdn5^40/40^*, *ΔPfdn5^15/40^*, and *actin5C-Gal4>UAS-Pfdn5; ∆Pfdn5^15/40^*. *rp49* was used as an internal control.

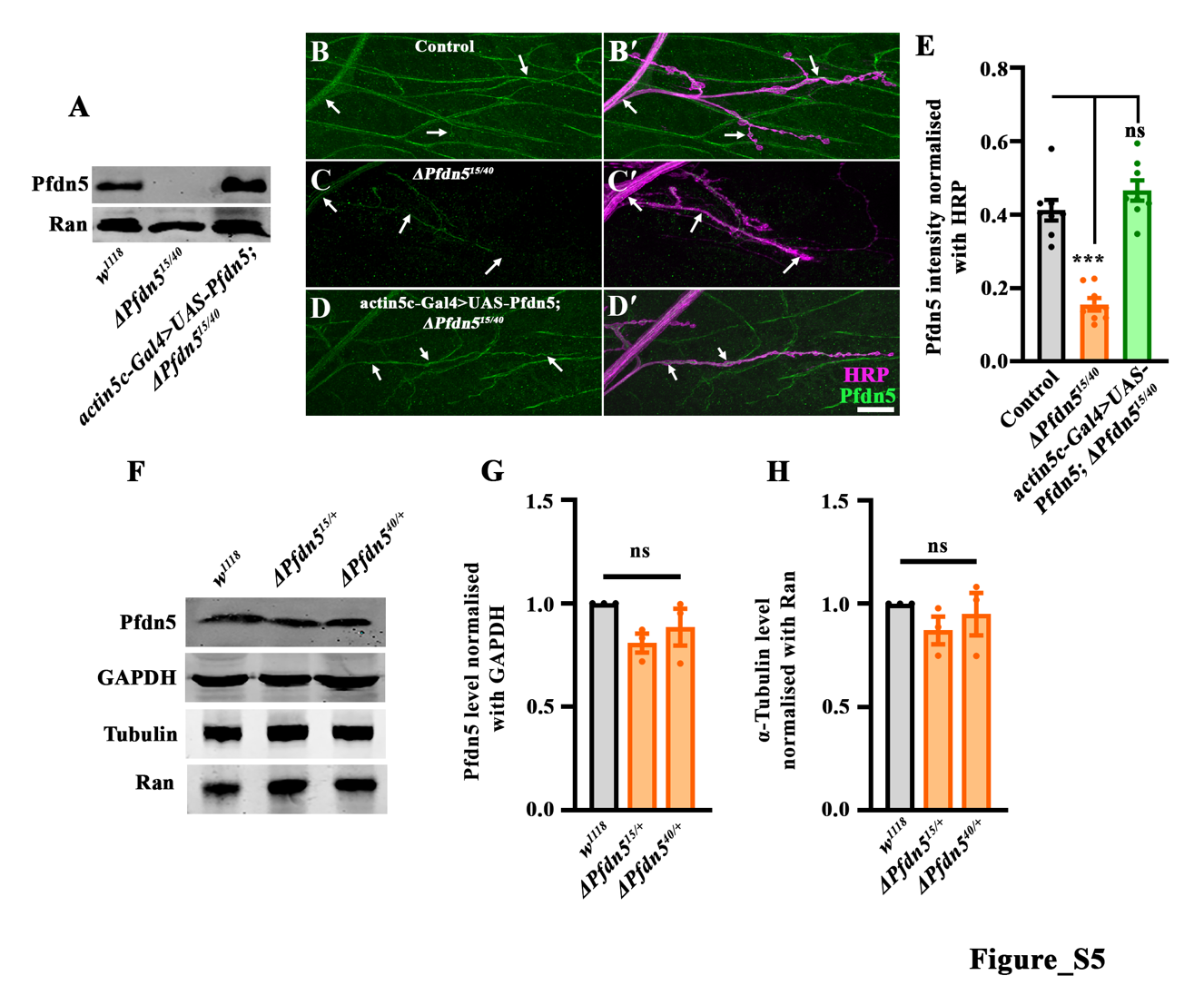

**Figure S5: Generation and characterization of Pfdn5 antibody**

**(A)** Western blot showing the levels of Pfdn5 in control, *ΔPfdn5^15/40,^* and *actin5c-Gal4>UAS-Pfdn5; ΔPfdn5^15/40^.* Ran was used as a loading control.

**(B-D')** Confocal images of NMJ synapses at muscle 4 of A2 hemisegment showing Pfdn5 levels in (C-C') control (D-D') *ΔPfdn5^15/40^* (E-E') *actin5c-Gal4>UAS-Pfdn5; ΔPfdn5^15/40^* double immunolabeled for HRP (magenta) and Pfdn5 (green). The scale bar in D' for (B-D') represents 10 µm.

**(E)** Histogram showing the average fluorescence intensity of Pfdn5 at the NMJ of muscle 4 in control (0.41 ± 0.02), *ΔPfdn5^15/40^* (0.15 ± 0.01), and *actin5c-Gal4>UAS-Pfdn5; ΔPfdn5^15/40^* (0.46 ± 0.02)*.* ***p<0.001; ns, not significant. At least 8 NMJs of each genotype were used for quantification.

**(F)** Western blot showing the levels of Pfdn5 and Tubulin in control, *ΔPfdn5^15/+,^* and *ΔPfdn5^40/+^.* GAPDH was used as a loading control for Pfdn5. Ran was used as a loading control for Tubulin.

**(G)** Histogram showing the level of Pfdn5 normalised with GAPDH in control (1.00 ± 0.00), *ΔPfdn5^15/+^* (0.81 ± 0.05), and *ΔPfdn5^40/+^* (0.89 ± 0.09)*.* ns, not significant.

**(H)** Histogram showing the level of Tubulin normalised with Ran in control (1.00 ± 0.00), *ΔPfdn5^15/+^* (0.87 ± 0.07), and *ΔPfdn5^40/+^* (0.95 ± 0.10)*.* ns, not significant.

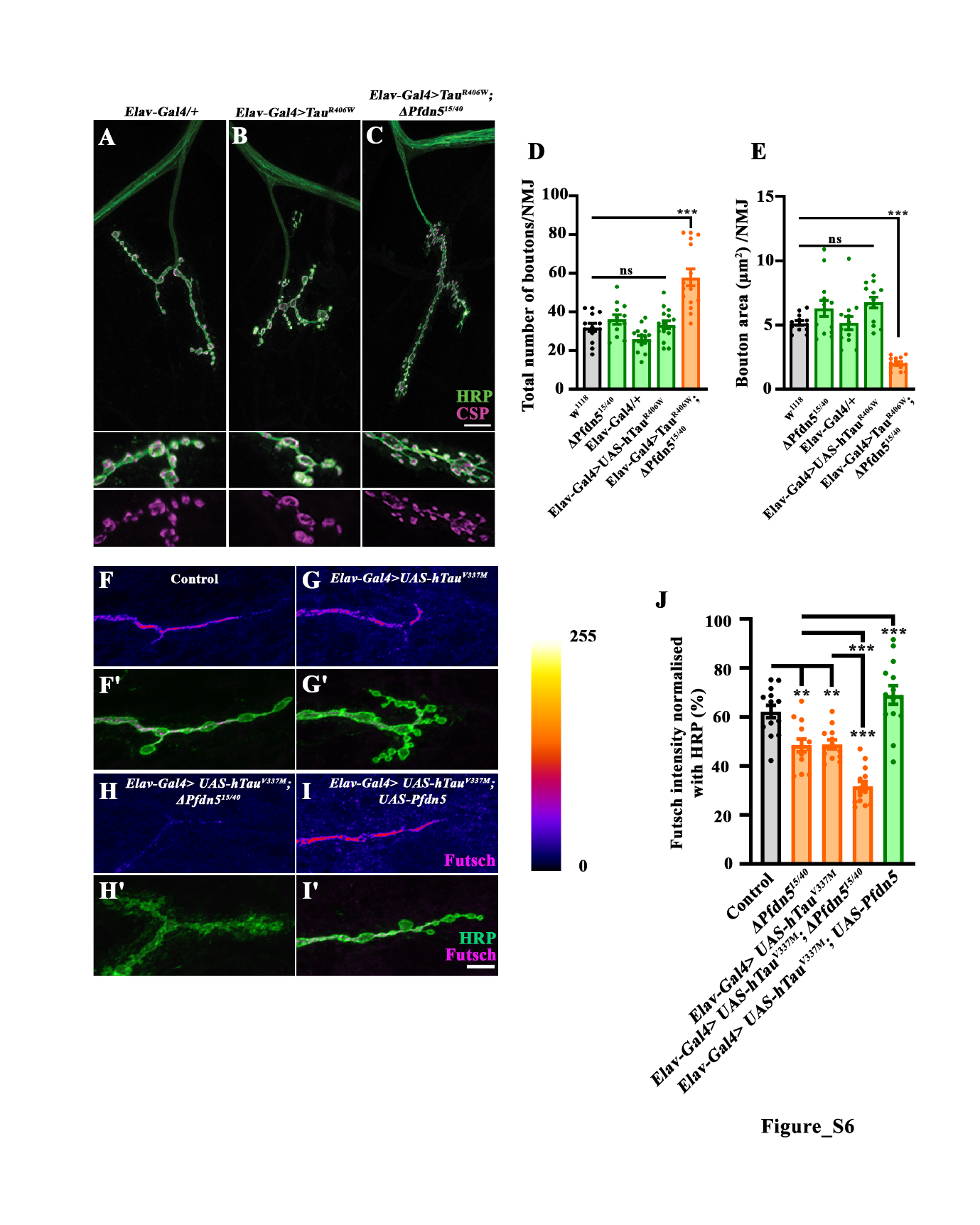

**Figure S6: Loss of Pfdn5 enhances hTau^R406W^-induced synaptic phenotypes**

**(A-C)** Confocal images of NMJ synapses at muscle 4 of A2 hemisegment showing synaptic morphology in control, *Elav-Gal4>hTau^R406W^,* and *Elav-Gal4>hTau^R406W^; ΔPfdn5^15/40^* double immunolabelled for CSP (magenta) and HRP (green). The inset shows the magnified images of the NMJs. The scale bar in the C represents 10 µm for images (A-C).

**(D)** Histogram showing total bouton number from muscle 4 at A2 hemisegment in control (32.08 ± 2.14), *ΔPfdn5^15/40^* (36.08 ± 2.50), *Elav-Gal/+* (26.13 ± 1.58), *Elav-Gal4>hTau^R406W^* (33.38 ± 1.97)*,* and *Elav-Gal4>hTau^R406W^; ΔPfdn5^15/40^* (57.87 ± 4.32). ***p<0.001; ns, not significant. At least 13 NMJs of each genotype were used for quantification.

**(E)** Histogram showing the average bouton area from muscle 4 at A2 hemisegment in control (5.15 ± 0.18), *ΔPfdn5^15/40^* (6.29 ± 0.62), *Elav-Gal/+* (5.15 ± 0.15), *Elav-Gal4>hTau^R406W^* (6.75 ± 0.43)*,* and *Elav-Gal4>hTau^R406W^; ΔPfdn5^15/40^* (2.0 ± 0.14). ***p<0.001; ns, not significant. At least 13 NMJs of each genotype were used for quantification.

**(F-I')** Confocal images of NMJ synapses at muscle 4 of A2 hemisegment showing futsch intensity in control (F-F'), *Elav-Gal4>hTau^V337M^* (G-G')*, Elav-Gal4>hTau^V337M^; ΔPfdn5^15/40^* (H-H') and *Elav-Gal4>hTau^V337M^; UAS-Pfdn5* (I-I') double immunolabelled for 22C10 (magenta) and HRP (green). The scale bar in the I' represents 10 µm for images (F-I').

**(J)** Histogram showing futsch intensity normalised with HRP from muscle 4 at A2 hemisegment in control (62.18 ± 2.56), *ΔPfdn5^15/40^* (48.32 ± 2.66), *Elav-Gal4>hTau^V337M^* (48.75 ± 1.79), *Elav-Gal4>hTau^V337M^; ΔPfdn5^15/40^* (31.58 ± 2.0)*,* and *Elav-Gal4>hTau^V337M^; UAS-Pfdn5* (68.94 ± 3.78). **p<0.01; ***p<0.001; ns, not significant. At least 13 NMJs of each genotype were used for quantification.

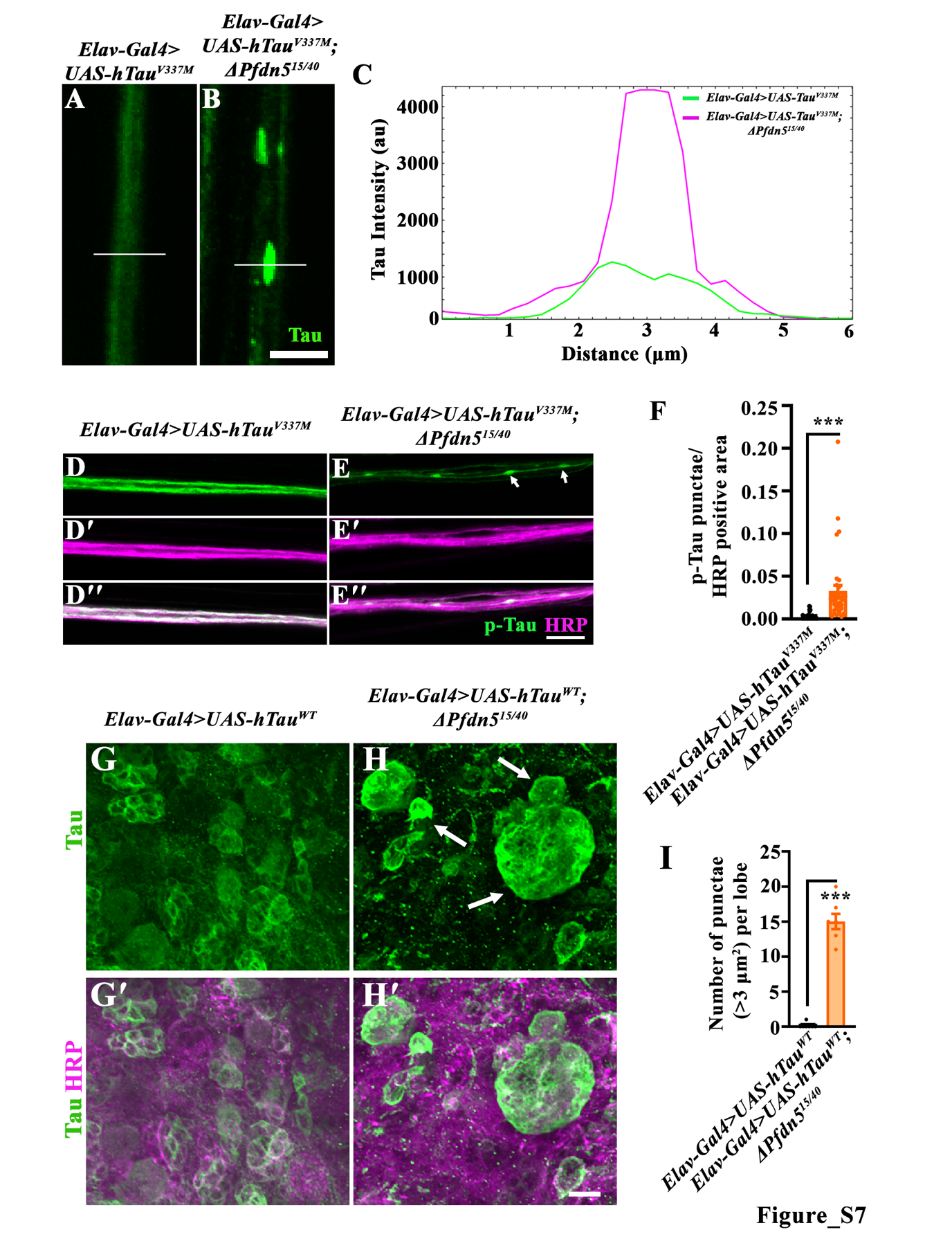

**Figure S7: hTau aggregates in the axons and larval brain of the *Pfdn5* mutants**

**(A-B)** Confocal images showing the intensity profile across the axons labeled with the T46 (total Tau) in *Elav-Gal4>hTau^V337M^* and *Elav-Gal4>hTau^V337M^*; *ΔPfdn5^15/40^*. The line drawn across the axons is 6 µm, and the scale bar in B represents 5 µm for (A-B).

**(C)** Intensity plot profile showing the intensity and distribution of the T46 across the axons in the *Elav-Gal4>hTau^V337M^* (green) and *Elav-Gal4>hTau^V337M^*; *ΔPfdn5^15/40^* (magenta)*.*

**(D-E'')** Confocal single-section images of third instar larval axons in (D-D'') *Elav-Gal4>UAS-hTau^V337M^*, (E-E'') *Elav-Gal4>UAS-hTau^V337M^; Pfdn5^15/40^* double immunolabeled with neuronal membrane marker, HRP (magenta), and AT8 antibody against phospho-Tau (green). The scale bar in E'' for (D-E'') represents 10 µm. Arrows in (E) point to the punctae or the aggregates of Tau.

**(F)** Histogram showing the quantification of the number of phospho-Tau punctae normalized with HRP positive area in *Elav-Gal4>UAS-hTau^V337M^* (0.002 ± 0.0005), *Elav-Gal4>UAS-hTau^V337M^; ΔPfdn5^15/40^* (0.03 ± 0.006). ***p<0.001; ns, not significant. At least 36 axons from 8 animals of each genotype were used for quantification.

**(G-H')** Confocal images of third instar larval brain showing hTau aggregates in *Elav-Gal4>UAS-hTau^WT^* (G-G'), *Elav-Gal4>UAS-hTau^WT^; Pfdn5^15/40^* (H-H') double immunolabeled with neuronal membrane marker, HRP (magenta), and T46 antibody against total Tau (green). The scale bar in H' for (G-H') represents 20 µm.

**(I)** Histogram showing the quantification of the number of total Tau punctae (>3 μm^2^) per lobe in *Elav-Gal4>UAS-hTau^WT^* (0.14 ± 0.14), *Elav-Gal4>UAS-hTau^WT^; ΔPfdn5^15/40^* (15 ± 1.09). ***p<0.001; ns, not significant. At least 7 lobes of each genotype were used for quantification.

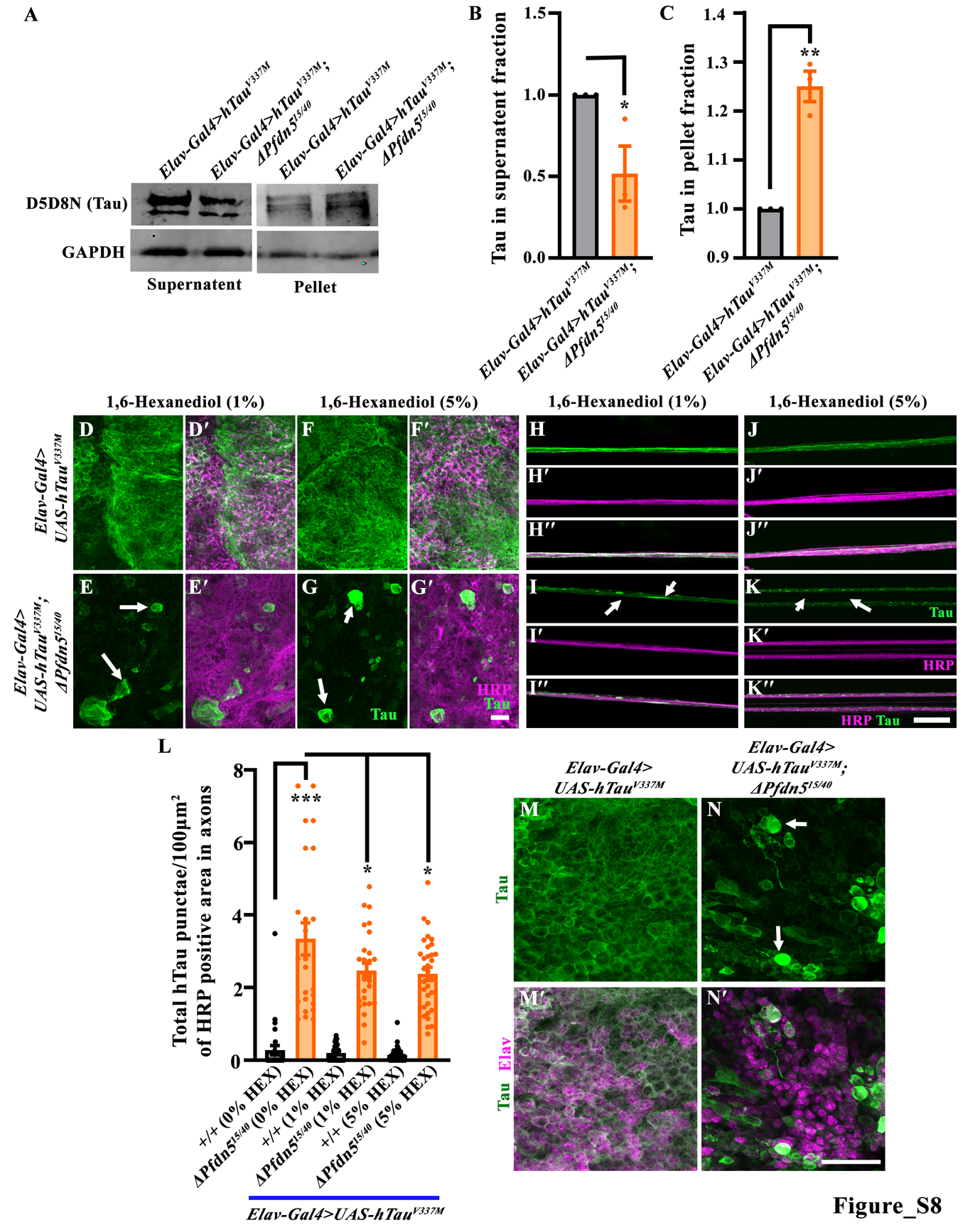

**Figure S8: Loss-of-Pfdn5 results in the formation of stable hTau aggregates**

**(A)** Western blot showing the Tau protein level in supernatant and pellet fraction of *Elav-Gal4>hTau^V337M^* and *Elav-Gal4>hTau^V337M^*; *ΔPfdn5^15/40^.* GAPDH was used as a loading control.

**(B)** Histogram showing the level of Tau in supernatant fraction normalised with GAPDH in *Elav-Gal4>hTau^V337M^* (1.00 ± 0.00), and *Elav-Gal4>hTau^V337M^*; *ΔPfdn5^15/40^* (0.52 ± 0.17). *p<0.05; ns, not significant.

**(C)** Histogram showing the level of Tau in pellet fraction normalised in *Elav-Gal4>hTau^V337M^* (1.00 ± 0.00), and *Elav-Gal4>hTau^V337M^*; *ΔPfdn5^15/40^* (1.25 ± 0.03). **p<0.01; ns, not significant.

**(D-G')** Confocal images of third instar larval brain showing hTau aggregates in *Elav-Gal4>hTau^V337M^* with 1% 1,6-Hexanediol (D-D'), *Elav-Gal4>hTau^V337M^*; *ΔPfdn5^15/40^* with 1% 1,6-Hexanediol (E-E'), *Elav-Gal4>hTau^V337M^* with 5% 1,6-Hexanediol (F-F'), *Elav-Gal4>hTau^V337M^*; *ΔPfdn5^15/40^* with 5% 1,6-Hexanediol (G-G') double immunolabeled with neuronal membrane marker, HRP (magenta), and T46 antibody against total Tau (green). The scale bar in G' for (D-G') represents 10 µm.

**(H-K'')** Confocal images of third instar larval axons showing hTau aggregates in *Elav-Gal4>hTau^V337M^* with 1% 1,6-Hexanediol (H-H''), *Elav-Gal4>hTau^V337M^*; *ΔPfdn5^15/40^* with 1% 1,6-Hexanediol (I-I''), *Elav-Gal4>hTau^V337M^* with 5% 1,6-Hexanediol (J-J''), *Elav-Gal4>hTau^V337M^*; *ΔPfdn5^15/40^* with 5% 1,6-Hexanediol (K-K'') double immunolabeled with neuronal membrane marker, HRP (magenta), and T46 antibody against total Tau (green). The scale bar in K'' for (H-K'') represents 10 µm.

**(L)** Histogram showing the quantification of the number of total Tau per 100 μm^2^ of HRP positive area in *Elav-Gal4>hTau^V337M^* with 0% 1,6-Hexanediol (0.28 ± 0.12), *Elav-Gal4>hTau^V337M^*; *ΔPfdn5^15/40^* with 0% 1,6-Hexanediol (3.35 ± 0.45), *Elav-Gal4>hTau^V337M^* with 1% 1,6-Hexanediol (0.21 ± 0.04), *Elav-Gal4>hTau^V337M^*; *ΔPfdn5^15/40^* with 1% 1,6-Hexanediol (2.47 ± 0.21), *Elav-Gal4>hTau^V337M^* with 5% 1,6-Hexanediol (0.15 ± 0.04), *Elav-Gal4>hTau^V337M^*; *ΔPfdn5^15/40^* with 5% 1,6-Hexanediol (2.39 ± 0.18). *p<0.05; ***p<0.001; ns, not significant. At least 24 axons from 8 animals of each genotype were used for quantification.

**(M-N')** Confocal images of third instar larval brain showing localisation of hTau aggregates in *Elav-Gal4>hTau^V337M^* (M-M'), *Elav-Gal4>hTau^V337M^*; *ΔPfdn5^15/40^* (N-N') double immunolabeled with Elav (magenta), and T46 antibody against total Tau (green). The scale bar in N' for (M-N') represents 10 µm.

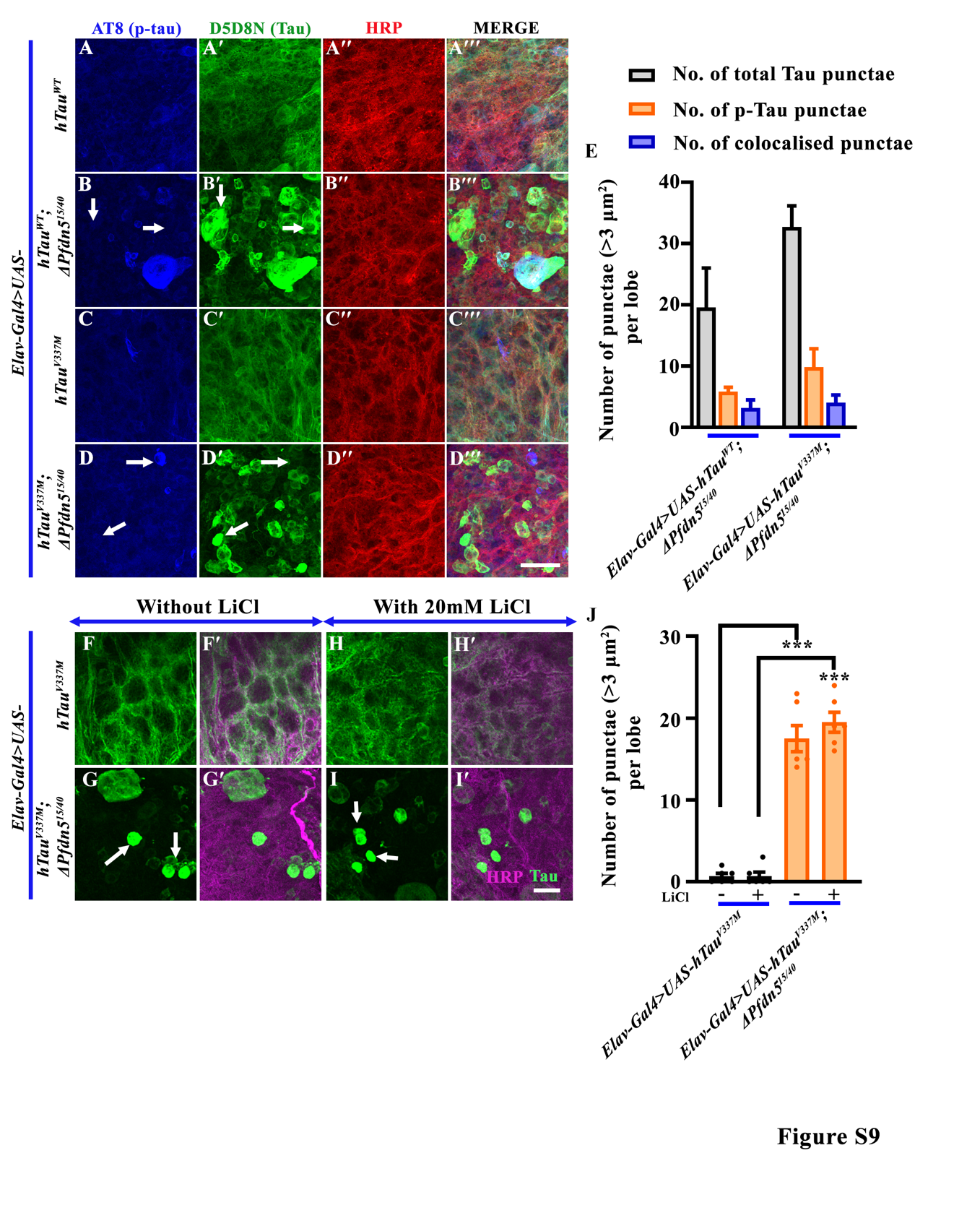

**Figure S9: Loss of Pfdn5 induces Tau-aggregation independent of hyperphosphorylation**

**(A-D''')** Confocal images of third instar larval brain showing hTau aggregates in *Elav-Gal4>hTau^WT^* (A-A'''), *Elav-Gal4>hTau^WT^*; *ΔPfdn5^15/40^* (B-B'''), *Elav-Gal4>hTau^V337M^* (C-C'''), *Elav-Gal4>hTau^V337M^*; *ΔPfdn5^15/40^* (D-D''') triple immunolabeled with neuronal membrane marker, HRP (red), AT8 antibody against p-Tau (blue) and D5D8N antibody against total Tau (green). The scale bar in D''' for (A-D''') represents 20 µm. The arrow points to the mutually exclusive Tau aggregates.

**(E)** Histogram showing the quantification of the number of Tau punctae (>3 μm^2^) per lobe in *Elav-Gal>UAS-hTau^WT^; ΔPfdn5^15/40^* and *Elav-Gal4>UAS-hTau^V337M^; ΔPfdn5^15/40^*.

**(F-I')** Confocal images of third instar larval brain showing hTau aggregates in *Elav-Gal4>hTau^V337M^* (F-F'), *Elav-Gal4>hTau^V337M^*; *ΔPfdn5^15/40^* (G-G') without LiCl, *Elav-Gal4>hTau^V337M^* (H-H'), *Elav-Gal4>hTau^V337M^*; *ΔPfdn5^15/40^* (I-I') in the presence of 20mM double immunolabeled with neuronal membrane marker, HRP (magenta), and T46 antibody against total Tau (green). The scale bar in I' for (F-I') represents 10 µm.

**(J)** Histogram showing the quantification of the number of total Tau punctae (>3 μm^2^) per lobe in *Elav-Gal4>hTau^V337M^* without LiCl (0.67 ± 0.33), *Elav-Gal4>hTau^V337M^* with LiCl (0.67 ± 0.49), *Elav-Gal4>hTau^V337M^*; *ΔPfdn5^15/40^* without LiCl (17.5 ± 1.61), *Elav-Gal4>hTau^V337M^*; *ΔPfdn5^15/40^* with LiCl (19.51 ± 1.32). ***p<0.001; ns, not significant. At least 6 lobes of each genotype were used for quantification.

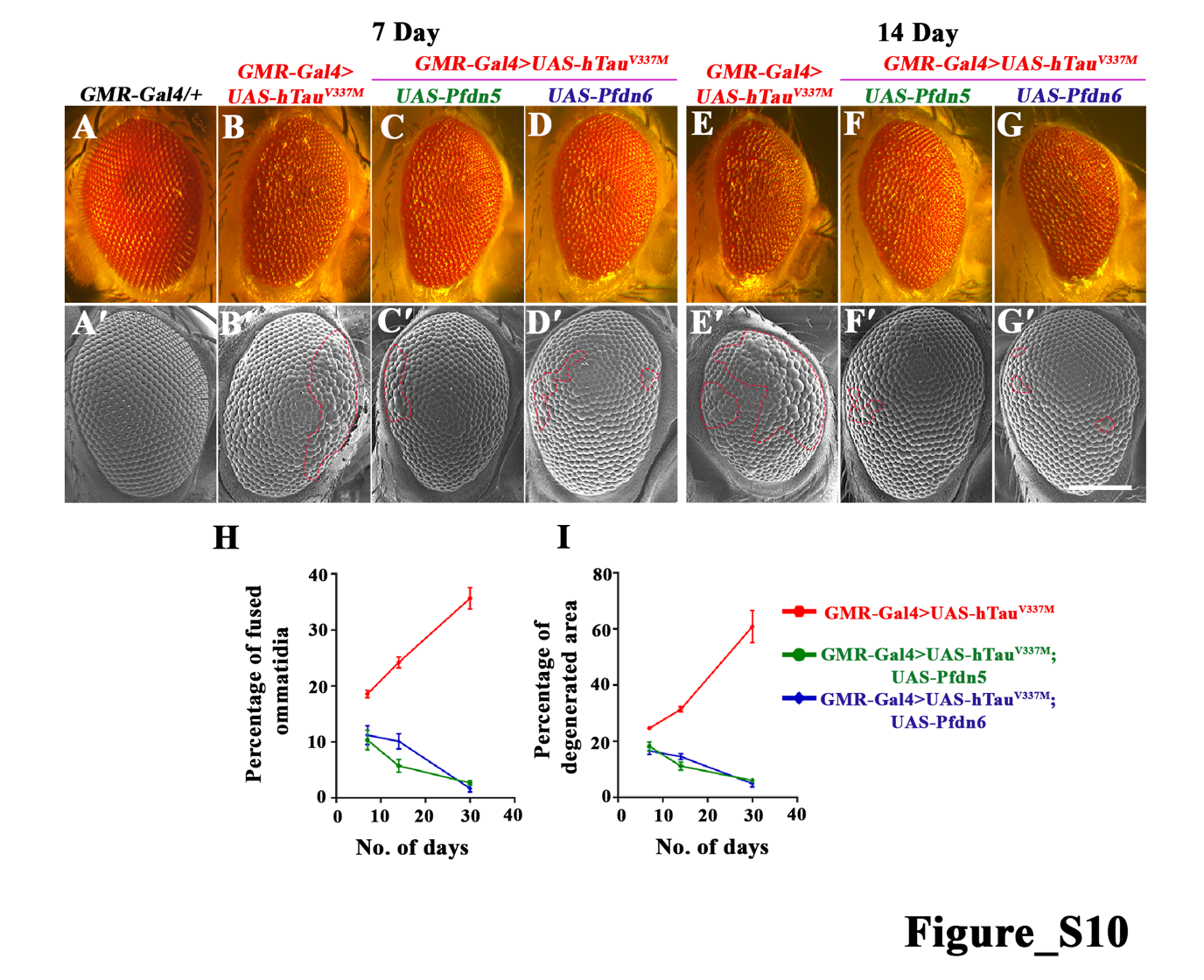

**Figure S10: Pfdn5 rescues progressive eye degeneration induced by expression of Tau^V337M^**

**(A-G)** Bright-field images of 7 and 14-day-old *Drosophila* eyes expressing (A) *GMR-Gal4/+* (control), (B and E) *GMR-Gal4>UAS-hTau^V337M^*, (C and F) *GMR-Gal4>UAS-hTau^V337M^; UAS-Pfdn5*, (D and G) *GMR-Gal4>UAS-hTau^V337M^; UAS-Pfdn6*.

**(A'-G')** Scanning electron microscopic images of 7 and 14-day-old *Drosophila* eyes expressing (A') *GMR-Gal4/+* (control), (B' and E') *GMR-Gal4>UAS-hTau^V337M^*, (C' and F') *GMR-Gal4>UAS-hTau^V337M^; UAS-Pfdn5*, (D' and G') *GMR-Gal4>UAS-hTau^V337M^; UAS-Pfdn6*. The scale bar in G' for (A-G') represents 100 µm.

**(H-I)** Graph showing quantifications of age-dependent progression of ommatidial fusion (H) and percentage of degenerated eye area (I) in *GMR-Gal4>UAS-hTau^V337M^*, *GMR-Gal4>UAS-hTau^V337M^; UAS-Pfdn5* and *GMR-Gal4>UAS-hTau^V337M^; UAS-Pfdn6*. Note that expression of Pfdn5 or Pfdn6 suppresses the Tau-induced progressive eye degeneration. At least 12 SEM eye images of each genotype were used for quantification at each time point.

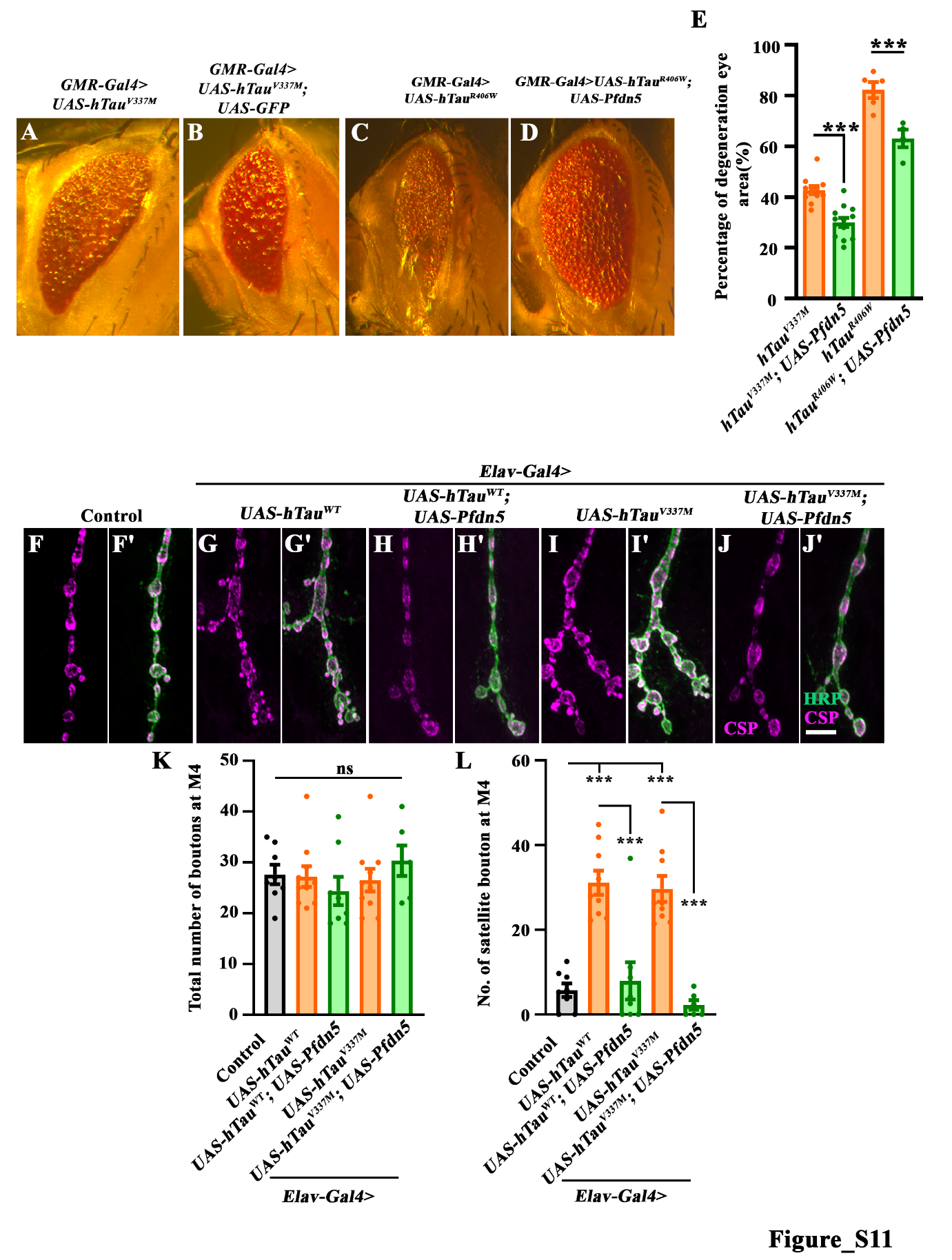

**Figure S11: Coexpression of Pfdn5 with hTau variants rescues the ommatidial degeneration and synaptic defects**

**(A-B)** Bright-field images of 30-day-old *Drosophila* eyes expressing (A) *GMR-Gal4>UAS-hTau^V337M^*, and (B) *GMR-Gal4>UAS-hTau^V337M^; UAS-GFP*.

**(C-D)** Bright-field images of 5-day-old *Drosophila* eyes expressing (E) *GMR-Gal4>UAS-hTau^R406W^*, and (F) *GMR-Gal4>UAS-hTau^R406W^; UAS-Pfdn5.*

**(E)** Histogram showing the percentage of degenerated area in *GMR-Gal4>UAS-hTau^V337M^* (42.74 ±7.6), *GMR-Gal4>UAS-hTau^V337M^; UAS-Pfdn5* (30.03 ± 1.74), *GMR-Gal4>UAS-hTau^R406W^* (82.15 ± 3.19), and *GMR-Gal4>UAS-hTau^R406W^; UAS-Pfdn5* (63.11 ± 3.49). At least 4 brightfield eye images of each genotype were used for quantification.

**(F-J')** Confocal images of NMJ synapses at muscle 4 of A2 hemisegment showing synaptic morphology in (F) *Elav-Gal4/+* (control), (G) *Elav-Gal4>UAS-hTau^WT^*, (H) *Elav-Gal4>hTau^WT^; UAS-Pfdn5*, (I) *Elav-Gal4>hTau^V337M^*, (J) *Elav-Gal4>hTau^V337M^; UAS-Pfdn5* double immunolabeled with Hoechst (cyan), and Phalloidin (magenta). Arrows in (F) point to the pathological vacuolar structures. The scale bar in J for (F-J') represents 10 µm.

**(K)** Histogram showing the number of boutons from muscle 4 at A2 hemisegment in *Elav-Gal4/+* (control) (G), *Elav-Gal4>UAS-hTau^WT^* (H), *Elav-Gal4>hTau^WT^; UAS-Pfdn5* (I), *Elav-Gal4>hTau^V337M^* (J), *Elav-Gal4>hTau^V337M^; UAS-Pfdn5.* ns; not significant. At least 16 NMJs of each genotype were used for quantification.

**(L)** Histogram showing the number of satellite boutons from muscle 4 at A2 hemisegment in *Elav-Gal4/+* (control) (G), *Elav-Gal4>UAS-hTau^WT^* (H), *Elav-Gal4>hTau^WT^; UAS-Pfdn5* (I), *Elav-Gal4>hTau^V337M^* (J), *Elav-Gal4>hTau^V337M^; UAS-Pfdn5.* ***p<0.001. At least 16 NMJs of each genotype were used for quantification.

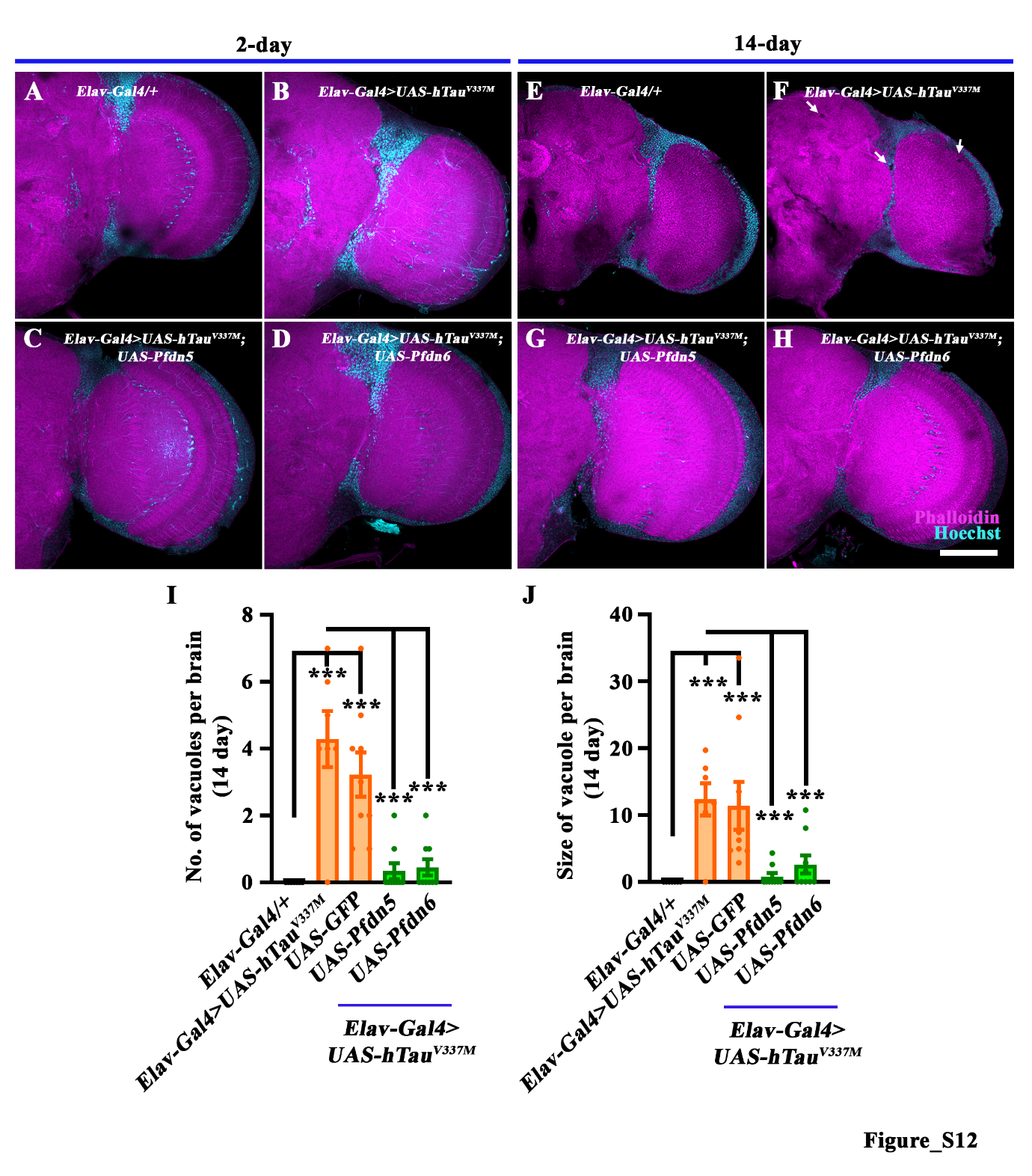
**Figure S12: Over-expression of Pfdn5 suppresses the age-dependent vacuolisation in hTau^V337M^-expressing flies.**

**(A-D)** Confocal images of a single section of a 2-day-old adult brain in (A) *Elav-Gal4/+* (control), (B) *Elav-Gal4>UAS-hTau^V337M^*, (C) *Elav-Gal4>hTau^V337M^; UAS-Pfdn5*, (D) *Elav-Gal4>hTau^V337M^; UAS-Pfdn6* double immunolabeled with Hoechst (cyan), and Phalloidin (magenta).

**(E-H)** Confocal images of a single section of 14-day-old adult brain in (E) *Elav-Gal4/+* (control), (F) *Elav-Gal4>UAS-hTau^V337M^*, (G) *Elav-Gal4>hTau^V337M^; UAS-Pfdn5*, (H) *Elav-Gal4>hTau^V337M^; UAS-Pfdn6* double immunolabeled with Hoechst (cyan), and Phalloidin (magenta). Arrows in (F) point to the pathological vacuolar structures. The scale bar in H for (A-D) represents 20 µm.

**(I)** Histogram showing the quantification of number of vacuoles in 14 days old adult brain in *Elav-Gal4/+* (0.00 ± 0.00), *Elav-Gal4>UAS-hTau^V337M^* (4.29 ± 0.83), *Elav-Gal4>hTau^V337M^; UAS-GFP* (3.22 ± 0.66), *Elav-Gal4>hTau^V337M^; UAS-Pfdn5* (0.33 ± 0.24), *Elav-Gal4>hTau^V337M^; UAS-Pfdn6* (0.44 ± 0.24). ***p<0.001. At least 7 brains of each genotype were used for quantification.

**(J)** Histogram showing the quantification of vacuole size (in µm^2^) in 14-day old adult brain in *Elav-Gal4/+* (0.00 ± 0.00), *Elav-Gal4>UAS-hTau^V337M^* (12.36 ± 2.40), *Elav-Gal4>hTau^V337M^; UAS-GFP* (11.39 ± 3.57), *Elav-Gal4>hTau^V337M^; UAS-Pfdn5* (0.77 ± 0.53), *Elav-Gal4>hTau^V337M^; UAS-Pfdn6* (2.58 ± 1.35). ***p<0.001. At least 7 brains of each genotype were used for quantification.

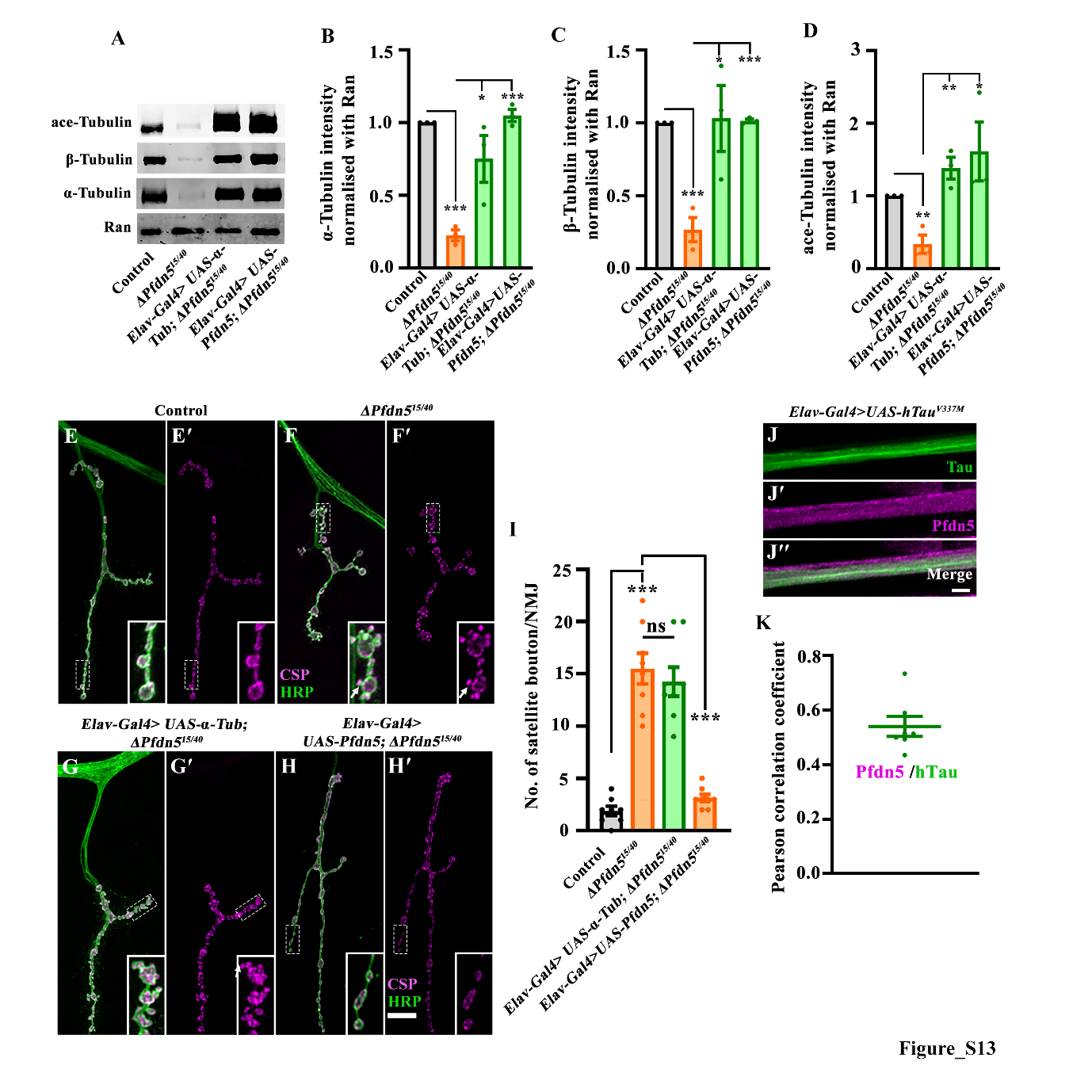

**Figure S13: Pfdn5 is required to stabilize microtubules at the synapses**

**(A)** Western blot showing protein levels of ace-Tubulin, α-Tubulin, and β-Tubulin in control, *∆Pfdn^15/40^,* *Elav-Gal4>UAS-α-Tubulin; ΔPfdn5^15/40^* and *Elav-Gal4>UAS-Pfdn5; ∆Pfdn5^15/40^*. Ran protein levels were used as an internal loading control.

**(B)** Histogram showing the percentage of α-Tubulin normalized with Ran in control (1.00 ± 0.00), *ΔPfdn5^15/40^* (0.23 ± 0.04), *Elav-Gal4>UAS-α-Tubulin; ΔPfdn5^15/40^* (0.75 ± 0.16), and *Elav-Gal4>UAS-Pfdn5; ∆Pfdn5^15/40^* (1.05 ± 0.04). ***p<0.001; *p<0.05. Three independent Western blots were used for quantification.

**(C)** Histogram showing the percentage of β-Tubulin normalized with Ran in control (1.00 ± 0.00), *ΔPfdn5^15/40^* (0.27 ± 0.08), *Elav-Gal4>UAS-α-Tubulin; ΔPfdn5^15/40^* (1.03 ± 0.23), and *Elav-Gal4>UAS-Pfdn5; ∆Pfdn5^15/40^* (1.01 ± 0.01). ***p<0.001; *p<0.05. Three independent Western blots were used for quantification.

**(D)** Histogram showing the percentage of ace-Tubulin normalized with Ran in control (1.00 ± 0.00), *ΔPfdn5^15/40^* (0.34 ± 0.13), *Elav-Gal4>UAS-α-Tubulin; ΔPfdn5^15/40^* (1.38 ± 0.15), and *Elav-Gal4>UAS-Pfdn5; ∆Pfdn5^15/40^* (1.61± 0.4). **p<0.01; *p<0.05. Three independent Western blots were used for quantification.

**(E-H')** Confocal images of NMJ synapses at muscle 4 of A2 hemisegment showing synaptic morphology in (E-E') control, (F-F') *ΔPfdn5^15/40^*, (G-G') *Elav-Gal4>UAS-α-Tubulin; ΔPfdn5^15/40^* (H-H') *Elav-Gal4>UAS-Pfdn5; ΔPfdn5^15/40^* double immunolabeled for HRP (green), and CSP (magenta). The scale bar in H for (E-H') represents 10 µm.

**(I)** Histogram showing number of satellite boutons from muscle 4 at A2 hemi segment in control (1.88 ± 0.44), *ΔPfdn5^15/40^* (15.5 ± 1.48), *Elav-Gal4>UAS-α-Tubulin; ΔPfdn5^15/40^* (14.25 ± 1.39), *Elav-Gal4>UAS-Pfdn5; ΔPfdn5^15/40^* (3.53 ± 0.32). ***p<0.001; ns, not significant. At least 8 NMJs of each genotype were used for quantification.

**(J-J')** Confocal images of third instar larval axons showing colocalization between Pfdn5 (magenta) and Tau (green). Scale bar in J' represents 10 µm.

**(K)** Pearson’s correlation coefficient to quantify colocalization between Pfdn5 and axonal microtubule.
