## Supplemental Table_1 for "Prefoldin 5 is a microtubule-associated protein that suppresses Tau-aggregation and neurotoxicity"

**Supplemental Table 1: The table shows list of HSPs used to screen as the genetic modifier of human Tau^V337M^. The values represent the percentage degenerated eye area.**

| **S. No.** | **Gene Family** | **CG number** | **Gene Name** | **VDRC/BDSC No.** | **Phenotype with GMR-Gal4** | **Phenotype with hTau^V337M^** | **Mean ± SEM** |
| --- | --- | --- | --- | --- | --- | --- | --- |
| 1 | Human Tau | hTau^V337M^ | Human Tau | - | Rough eye | Control | 41.69 ± 3.51 |
| 2 | Prefoldin | CG10635 | Prefoldin 4 | BL 77412 | No | Enhancement | 63.58 ± 2.95 |
| 3 |  | CG7048 | Prefoldin 5 | BL 67815 | No | Enhancement | 61.56 ± 3.62 |
|  |  |  |  | KK100796 | No | Enhancement | 85.06 ± 3.01 |
|  |  |  |  | GD29812 | No | Enhancement | 64.44 ± 4.92 |
| 4 |  | CG7770 | Prefoldin 6 | BL65365 | No | Enhancement | 70.95 ± 4.17 |
|  |  |  |  | GD34204 | Glossy eye | Enhancement | 98.57 ± 0.36 |
|  |  |  |  | KK101541 | No | Enhancement | 62.01 ± 5.69 |
| **5** |  | CG6302 | Prefoldin 2 | BL-61843 | No | No change | - |
| **6** |  | CG13993 | Prefoldin 1 | BL66998 | No | No change | - |
| **7** | Tbce -Tubulin-binding cofactor E | CG7861 | TBCE | BL-34537 | No | Enhancement | 65.3 ± 3.65 |
|  |  |  |  | BL-44569 | No | No change | - |
| **8** | Chaperonin containing TCP1 complex | CG8439 | CCT5 | BL-41818 | No | Enhancement | 64.36 ± 2.49 |
|  |  |  |  | BL-57440 | No | No change | - |
|  |  |  |  | KK109505 | No | No change | - |
| **9** |  | CG5525 | CCT4 | KK106099 | No | No change |  |
|  |  |  |  | GD22155 | No | No change |  |
| **10** |  | CG8351 | CCT7 | BL-34931 | No | Enhancement | 61.67 ± 4.47 |
|  |  |  |  | KK108585 | Glossy eye | Enhancement | 61.00 ± 5.67 |
| **11** |  | CG8977 | CCT3 | BL-34969 | No | No Change | - |
|  |  |  |  | GD36070 | No | Enhancement | 93.16 ± 1.24 |
| **12** |  | CG8231 | CCT6 | KK109734 | No | No Change |  |
| **13** |  | CG8258 | CCT8 | GD45789 | No | No Change | - |
| **14** | HSP100 | CG4538 | ClpX - Caseinolytic protease chaperone subunit | GD39699 | No | Enhancement | 74.67 ± 6.71 |
|  |  |  |  | GD16432 | No | Enhancement | 55.33 ± 1.44 |
| **15** | HSP90 | CG1242 | Heat shock protein 83 | KK108568 | Mildly rough eye | Enhancement | 65.02 ± 5.76 |
| **16** | HSP70 | CG8542 | Heat shock protein 70 cognate 5 | KK106236 | No | No change | - |
|  |  |  |  | GD47745 | No | Enhancement | 46.18 ± 3.84 |
| **17** |  | CG7756 | Heat shock protein 70 cognate 2 | BL-44485 | No | Enhancement | 77.32 ± 6.59 |
|  |  |  |  | BL-42014 | No | No change |  |
|  |  |  |  | GD19202 | No | Enhancement | 89.27 ± 1.98 |
| **18** |  | CG31366 | Heat shock protein 70 Aa | GD41748 | Rough eye | Enhancement | 90.62 ± 0.96 |
| **19** |  | CG4264 | Heat shock protein 70 cognate 4 | BL28709 | No | No change |  |
|  |  |  |  | BL-35684 | No | No change |  |
|  |  |  |  | GD26465 | Mild | Enhancement | 69.68 ± 5.98 |
| **20** |  | CG4147 | Heat shock protein 70 cognate 3 | BL-80420 | **LETHAL** | **LETHAL** | - |
|  |  |  |  | KK101766 | **LETHAL** | **LETHAL** | - |
|  |  |  |  | GD14882 | **LETHAL** | **LETHAL** | - |
| **21** |  | CG31449 | Heat shock protein 70 Ba | BL43289 | No | No change | - |
|  |  |  |  | BL-35672 | No | No change | - |
|  |  |  |  | GD50382 | **LETHAL** | **LETHAL** | - |
| **22** |  | CG5834 | Heat shock protein 70 Bbb | BL33000 | No | No change | - |
| **23** |  | CG8937 | Heat shock protein 70 cognate 1 | BL-34527 | No | No change | - |
|  |  |  |  | KK106510 | No | Suppression | 21.59 ± 1.28 |
| **24** |  | CG6489 | Heat shock protein 70 Bc | BL-42626 | No | Suppression | 13.83 ± 1.26 |
| **25** |  | CG31366, CG18743 | Hsp70Aa, Hsp70Ab | BL42639 | No | Suppression | 14.69 ± 2.30 |
|  |  |  |  | BL-35663 | No | Suppression | 3.89 ± 1.74 |
| **26** |  | CG7756, CG31449, CG31359, CG5834, CG6489 | Hsc70-2, Hsp70Ba, Hsp70Bb, Hsp70Bbb, Hsp70Bc | BL-32997 | No | No change | - |
| **27** |  | CG31449, CG31359, CG5834, CG6489 | Hsp70Ba, Hsp70Bb, Hsp70Bbb, Hsp70Bc | BL-28787 | No | No change | - |
| **28** | HSP60 chaperonins- Group 1 | CG12101 | Heat shock protein 60A | BL-34729 | No | No Change | - |
|  |  |  |  | GD18738 | No | Enhancement | 68.89 ± 3.09 |
| **29** |  | CG2830 | Heat shock protein 60B | BL-66328 | No | No Change | - |
| **30** |  | CG7235 | Heat shock protein 60C | BL67003 | No | No Change | - |
| **31** |  | CG16954 | Heat shock protein 60D | BL-57252 | No | No Change | - |
| **32** | HSP40 | CG1107 | auxilin | BL-39017 | No | Suppression | 27.32 ± 2.29 |
|  |  |  |  | KK103426 | No | No change | - |
| **33** |  | CG8863 | DnaJ-like-2 | BL-57382 | No | Enhancement | 65.47 ± 6.04 |
|  |  |  |  | KK104880 | No | Slight enhancement | - |
| **34** |  | CG8448 | mrj | BL66921 | No | No change | - |
|  |  |  |  | KK109817 | Glossy eye | Enhancement | 54.22 ± 2.71 |
| **35** |  | CG30156 | - | BL77162 | No | No change | - |
|  |  |  |  | BL67920 | No | Enhancement | 68.75 ± 4.39 |
|  |  |  |  | GD2714 | No | Enhancement | 65.07 ± 4.46 |
| **36** |  | CG9089 | wurst | BL58265 | No | No change | - |
|  |  |  |  | KK110270 | Glossy and rough eye | Enhancement | 70.24 ± 1.84 |
| **37** |  | CG5504 | - | BL-28594 | No | Enhancement | 65.56 ± 1.92 |
| **38** |  | CG6395 | Cysteine string protein | BL-31290 | No | Enhancement | 59.74 ± 3.62 |
| **39** |  | CG10578 | DnaJ-like-1 | BL-32978 | No | Suppression | 31.39 ± 1.09 |
|  |  |  |  | BL-32899 | No | No change | - |
|  |  |  |  | KK104618 | No | No change | - |
|  |  |  |  | GD31271 | No | No change | - |
| **40** |  | CG7133 | - | BL42820 | No | Suppression | 21.18 ± 2.99 |
|  |  |  |  | BL-60459 | No | No change | - |
| **41** |  | CG7130 |  | BL57854 | No | Suppression | 26.04 ± 1.51 |
| **42** |  | CG4164 | shriveled | BL-37507 | No | Suppression | 22.56 ± 1.67 |
|  |  |  |  | BL-54797 | No | No change | - |
| **43** |  | CG10565 | - | BL-62205 | No | No change | - |
|  |  |  |  | BL-43205 | No | Suppression | 21.12 ± 1.77 |
| **44** |  | CG1409 |  | BL-58200 | No | Suppression | 11.98 ± 1.90 |
| **45** |  | CG4599 | Tpr2 - Tetratricopeptide repeat protein 2 | BL-80386 | No | Suppression | 27.00 ± 0.76 |
|  |  |  |  | BL-53259 | No | No change | - |
| **46** |  | CG6693 | - | BL-35699 | No | No change | - |
|  |  |  |  | BL-43190 | No | No change | - |
| **47** |  | CG2790 | - | BL-58341 | No | No change | - |
|  |  |  |  | BL-67270 | No | No change | - |
| **48** |  | CG12020 | - | BL-62882 | No | No change | - |
| **49** |  | CG5268 | black pearl | BL-67828 | No | No change | - |
| **50** |  | CG10375 | - | BL-67938 | No | No change | - |
| **51** |  | CG14650 | - | BL-61972 | No | No change | - |
| **51** |  | CG2887 | - | BL-66983 | No | No change | - |
| **53** |  | CG2239 | jdp | BL67329 | No | No change (black pigmentation) | - |
| **54** |  | CG9828 | DnaJ homolog | BL61931 | No | No change | - |
| **55** |  | CG7556 | - | BL32511 | No | No change | - |
|  |  |  |  | KK107020 | Glossy and rough eye | Enhancement | 60.59 ± 3.29 |
| **56** |  | CG8014 | Rme-8 - Receptor mediated endocytosis 8 | KK107706 | No | No change | - |
| **57** |  | CG7394 |  | KK101490 | No | No change | - |
| **58** |  | CG40178 | lethal (3) 80Fg | BL-44578 | No | No change | - |
|  |  |  |  | KK109162 | No | No change | - |
|  |  |  |  | KK110089 | No | No change | - |
| **59** |  | CG8583 | Secretory 63 | BL52925 | No | No change | - |
|  |  |  |  | KK110331 | No | No change | - |
|  |  |  |  | GD33281 | No | No change | - |
| **60** |  | CG17187 | CWC23 | BL64480 | No | No change | - |
|  |  |  |  | BL55702 | No | No change | - |
| **61** |  | CG3061 | - | BL-66343 | No | No change | - |
| **62** |  | CG32641, CG32640 | - | BL65200 | No | No change | - |
| **63** | sHSPs | CG4461 | - | KK100857 | No | Suppression | 28.07 ± 3.19 |
| **64** |  | CG4463 | Heat shock protein 23 | KK102493 | No | Suppression | 22.92 ± 4.49 |
| **65** |  | CG4466 | Heat shock protein 27 | KK101669 | No | No change | - |
