## Supplemental Table_2 for "Prefoldin 5 is a microtubule-associated protein that suppresses Tau-aggregation and neurotoxicity"

Supplemental Table 2: List of primers used in this study

| Primer details | |
| --- | --- |
| gRNA1FP | 5′-TATATAGGAAAGATATCCGGGTGAACTTCGAGCAGGTGCGTTCCGCATGGTTTTAGAGCTAGAAATAGCAAG-3′ |
| gRNA1RP | 5′- ATTTTAACTTGCTATTTCTAGCTCTAAAACTAGTCTAGGAGCTCTGGGTACGACGTTAAATTGAAAATAGGTC-3′ |
| gRNA2FP | 5′- TATATAGGAAAGATATCCGGGTGAACTTCGTTAGGAGATTACCAACGTACGTTTTAGAGCTAGAAATAGCAAG-3′ |
| gRNA2RP | 5′- ATTTTAACTTGCTATTTCTAGCTCTAAAACGCACGTACTCCACGCGTCGCGACGTTAAATTGAAAATAGGTC-3′ |
| Pfdn5_FP1 | 5'- CGCATACAAATACAGAATGACTGAC-3' |
| Pfdn5_RP | 5'- CAAGACAAGTGTGTGCTC-3' |
| Pfdn5_Seq FP | 5'-GACACAGCGCGTACGTCCTTCG-3' |
| Pfdn5_RTFP | 5'-AGTCCGAGATGATCGACCTG-3' |
| Pfdn5_RTRP | 5'-CACCAGGATCTGACGGTTCT-3' |
| RP49_FP | 5'-AGATCGTGAAGAAGCGCACC-3' |
| RP49_RP | 5'- CGATCCGTAACCGATGTTGG-3' |
| Pfdn5_pET28 FP | 5′-ATGGCTGCCACCCCAAGCGCACC-3′ |
| Pfdn5_pET RP | 5′-GCCGCCAATGGGTCTCGAGGATC-3′ |
| Pfdn6 qRT FP | 5′-AAAGATGCAGGCCGAGATT-3′ |
| Pfdn6 qRT RP | 5′-ACGCACTTGTTCTCGTTCA-3′ |
| Pfdn5 qRT FP | 5′-CCTGACCAGCAGTATGTATGTG-3′ |
| Pfdn5 qRT RP | 5′-GTCGCTTGAAGTAGTCTTTGGA-3′ |
| Pfdn4 qRT FP | 5′-TCAAAGCGGAGCTGGAAA-3′ |
| Pfdn4 qRT RP | 5′-CCAACCAGGAACGGTATGT-3′ |
| TBCE qRT FP | 5′-GCAGAAGTCGATCGAACAAAGA-3′ |
| TBCE qRT RP | 5′-AGTAAGCGCAAAGTCGTGAG-3′ |
